# Effector-Specific Neural Representations of Perceptual Decisions Independent of Motor Actions and Sensory Modalities

**DOI:** 10.1101/2024.07.23.604761

**Authors:** Marlon F. Esmeyer, Timo T. Schmidt, Felix Blankenburg

## Abstract

Neuroscientific research has shown that perceptual decision-making occurs in effector-specific brain regions that are associated with the required motor response. Recent functional magnetic resonance imaging (fMRI) studies that dissociated decisions from coinciding processes, such as motor actions partly challenge this, indicating abstract representations that might vary across stimulus modalities. However, cross-modal comparisons have been difficult since most task designs differ not only in modality but also in effectors, motor response, and level of abstraction. Here, we describe an fMRI experiment where participants compared frequencies of two sequentially presented visual flicker stimuli in a delayed match-to-comparison task, which controlled for motor actions and stimulus sequence. Using Bayesian modelling, we estimated subjective frequency differences based on the time order effect. These values were applied in support vector regression analysis of a multi-voxel pattern whole-brain searchlight approach to identify brain regions containing information on subjective decision values. Furthermore, a conjunction analysis with data from a re-analyzed analogue vibrotactile study was conducted for a cross-modal comparison. Both analyses revealed significant activation patterns in the left dorsal (PMd) and ventral (PMv) premotor cortex as well as in the bilateral intraparietal sulcus (IPS). While previous primate and human imaging research have implicated these regions in transforming sensory information into action, our findings indicate that the IPS processes abstract decision signals while PMd and PMv represent an effector-specific, but motor response independent encoding of perceptual decisions that persists across sensory domains.

## Introduction

Humans rely on a variety of sensory information to navigate the multitude of every day’s decisions. Thus, understanding the neural mechanisms of perceptual decision-making has been a fundamental inquiry of neuroscientific research that has been explored across various sensory modalities using diverse research paradigms. An important avenue of research has investigated the neural correlates of vibrotactile perceptual decisions in primates (LaMotte & Mountcastle, 1975; Romo & de Lafuente, 2013). These studies typically employ variants of delayed match-to-comparison (DMTC) tasks, where participants compare frequencies of two subsequently presented vibrotactile flutter stimuli and decide whether the frequency of the second stimulus (f2) was higher or lower than the frequency of the first stimulus (f1). The response is usually indicated with a button press. In their seminal work, Romo and colleagues have described perceptual decision-making in DMTC tasks as a sequence of processing steps that dynamically link action and perception. Following the initial encoding of f1 in somatosensory areas, information on stimulus frequencies is retained across different neurons in higher somatosensory and frontal areas (see below), showing opposite tuning curves for high and low frequencies. This mnemonic representation of f1 is maintained during the delay-period until the presentation of f2. Subsequently, as f2 is encoded in the same areas, the decision whether f2 was higher or lower than f1 likely arises through the generation of a difference signal between neurons whose firing can be described by opposite tuning curves (Romo & de Lafuente, 2013). Following this computation, which corresponds to a subtraction of the two frequencies, a binary signed decision signal emerges, which is then transformed into a motor command, observed within the primary motor cortex. Notably, alongside prefrontal (Jun et al., 2010) and secondary somatosensory regions (Romo et al., 2002), decision-related signals have been most prominently observed across different effector-specific brain areas. In the context of the vibrotactile DMTC task, these encompass motor-related areas involved in the planning and execution of button presses or arm movements. Accordingly studies have reported decision-related activity in different sites of the premotor cortex, including the medial premotor cortex (PMm; de Lafuente & Romo, 2005; Hernández et al., 2002), the dorsal premotor cortex (PMd; Haegens et al., 2011; Rossi-Pool et al., 2017) and the ventral premotor cortex (PMv; Romo et al., 2004).

The findings by Romo and colleagues are in line with the intentional framework, which suggests that perceptual decisions are processed in brain areas associated with the required motor response (Shadlen et al., 2008). Similar to Romo and colleagues’ primate studies, human electroencephalography (EEG) studies using similar vibrotactile DMTC tasks revealed that beta band amplitude in the premotor cortex (Herding et al., 2016) and parietal event related potentials (Herding et al., 2019) were predictive of categorical choices. However, in most decision-making tasks, decisions are inextricably linked to task components such as the response, making it difficult to pinpoint neural markers of perceptual decisions independent of other sensory or motor processes. Human imaging studies that decoupled decision-from motor-related signals by employing multiple response-modalities or flexible response-mappings reported decision-related signals in varying brain region including abstract (Filimon et al., 2013; Heekeren et al., 2006), stimulus-modality specific (Hebart et al., 2012; Liu & Pleskac, 2011) or effector-specific representations (Hebart et al., 2016; Wu et al., 2019). These findings suggest that if a decision is mapped to an abstract decision-rule, it is represented in higher-order brain regions, while a decision that is directly mapped to a response is encoded in effector-specific brain regions. This notion was supported by an EEG study, which showed that beta band power encoded decisions in the premotor cortex when the motor response was known and in the parietal cortex when the decision was decoupled from the motor response (Ludwig et al., 2018). Interestingly, two recent vibrotactile DMTC studies that decoupled decisions from the response and the stimulus order in an functional magnetic resonance imaging (fMRI) experiment nevertheless decoded categorical choices from effector-specific brain regions, i.e. frontal eye fields (FEF) for saccadic responses, PMd for button presses and the intraparietal sulcus (IPS) for both (Wu et al., 2019, 2021). This indicates that effector-specific regions do not process perceptual decisions as concrete motor plans but instead in a more abstract, motor response independent manner, if a response effector is pre-specified.

To characterize the basic neural mechanisms underlying perceptual decision-making it is crucial to examine the similarities and differences across sensory modalities. Most of the perceptual learning literature has associated improvements in the perception of sensory stimuli with processes in early sensory cortices linked to specific low-level stimuli features that interact with feedback from higher-order brain regions (Fahle, 2005; Gilbert et al., 2001). This indicates potential fine-grained differences across stimulus modalities that might also apply to the closely associated perceptual decision processes. However, some studies investigating visual perceptual learning in the context of decision tasks have suggested that visual perceptual learning is at least partly effector-specific, with the posterior parietal cortex (PPC) as a likely candidate region (Ivanov et al., 2024; Law & Gold, 2008), suggesting effector-specific mechanisms for perceptual decisions that potentially persists across sensory modalities. Notably, comparing perceptual decisions has been challenging due to distinct paradigms used to investigate perceptual decisions in different modalities, most notably differing in the applied motor responses. For instance, while vibrotactile decisions have been investigated using DMTC paradigms with button presses as the response modality, research on visual decisions has typically employed random dot motion tasks and saccades as the response effector (Gold & Shadlen, 2007; Shadlen et al., 1996). Although earlier visual studies identified decision-related signals from neurons in effector-specific regions associated with planning and preparation of saccades, notably in the FEF and lateral intraparietal (LIP) neurons (Ding & Gold, 2012; Roitman & Shadlen, 2002; Shadlen & Newsome, 2001), newer findings suggest that these neurons represent aspects of decisions that occur independent of the effector modality (Shushruth et al., 2022; So & Shadlen, 2022; Zhou et al., 2023). Furthermore, a human neuroimaging study that decoupled decisions from motor responses in a visual random dot motion task nevertheless reported decision-related signals in the IPS (the human counterpart to LIP) and in the FEF (Liu & Pleskac, 2011). This indicates that IPS and FEF encode decisions according to the stimulus-modality (i.e. visual motion direction) rather than to the effector (i.e. eye movement). In contrast, the abovementioned vibrotactile studies found decision-related activation patterns in effector-specific brain regions, i.e. FEF for saccades and PMd for button presses as well as in the IPS (Wu et al., 2019, 2021). Considering these distinct findings, one can conclude that visual decisions might be represented stimulus-modality-specific and vibrotactile decisions in an effector-specific framework. However, unlike Liu & Pleskac (2011), in the studies by Wu et al. (2019, 2021) participants knew the effector modality they would use to select a response at the time of the decision. Thus, the observed differences might stem from variations in task paradigms rather than solely from differing processes between the sensory modalities of the task. Therefore, it is crucial to investigate cross-modal differences in a controlled design to identify underlying similarities and differences in neural processing.

Additional to task-related constrains, decision-making can be influenced by cognitive processes, such as the time-order effect (TOE), also known as contraction bias (Ashourian & Loewenstein, 2011; Herding et al., 2019). The TOE describes the phenomenon that the magnitude of a stimulus maintained over a delay period will be drawn towards the mean of previously observed stimuli (Boboeva et al., 2024). In DMTC tasks, this results in participants overestimating small f1 frequencies and underestimating large ones, which results in distinct performance distributions (Karim et al., 2012). To account for effects of the TOE, Herding et al. (2016, 2019) used a variant of the Bayesian model by Ashourian & Loewenstein (2011) to estimate subjective decision values in a DMTC task, suggesting that EEG signatures in premotor and parietal regions were modulated by subjective decision values rather than by the true underlying frequency differences. Thus, utilizing the TOE to infer on a continuous estimations of a subjective decision variable rather than the observed binary choice outcome is a promising approach to obtain more precise neural correlates of perceptual decisions.

The aim of the present fMRI study is to investigate cross-modal, i.e. visual and vibrotactile, neural correlates of perceptual decisions, which are independent of motor responses and stimulus sequence. Therefore, we used a visual adaptation of the vibrotactile DMTC task of Wu et al., (2021). To identify brain regions which process subjective decision values, we applied a multi-voxel pattern analysis (MVPA) whole-brain searchlight approach (Kriegeskorte et al., 2006) using support vector regression (SVR). To this end, we utilized subjective frequency differences (SFDs) estimated by a Bayesian model to account for the TOE. Furthermore, we reanalysed the data from Wu et al., (2021) using the same Bayesian modelling and SVR pipeline. We then conducted a conjunction analysis of both datasets to reveal how neural activation patterns reflect subjective decision values across sensory modalities. We hypothesized effector-specific activation patterns of subjective decision values in both the left PMd and the left IPS. Moreover, we expected these regions to predict subjective decision values consistently across the visual and the vibrotactile domain.

## Methods

### Participants

A total of 36 healthy volunteers (18 males, 18 females) with a mean age of 25.5 (standard deviation [SD] = 3.71, range: 18-34) participated in the fMRI experiment. Eligible participants were required to be right-handed, as assessed by the Edinburgh Handedness Inventory (Oldfield, 1971; 0.84 +-0.17) and free from any neurological or psychiatric disorders. All participants provided written informed consent and were compensated monetarily for their participation (12 Euros/h). The study was approved by the local ethics committee of the Freie Universität Berlin (003/2021).

### Task Design and Stimuli

The task design was adapted from the vibrotactile DMTC task by Wu et al., (2021). In our visual version of the DMTC task, we instructed participants to compare the frequency of two sequentially presented visual flicker stimuli (Fig. 1). F1 was 16, 20, 24 or 28 Hz and f2 was either 4 Hz above or below the first stimulus frequency (i.e. eight different frequency combinations in the frequency range of 12-32 Hz).

**Fig. 1.**
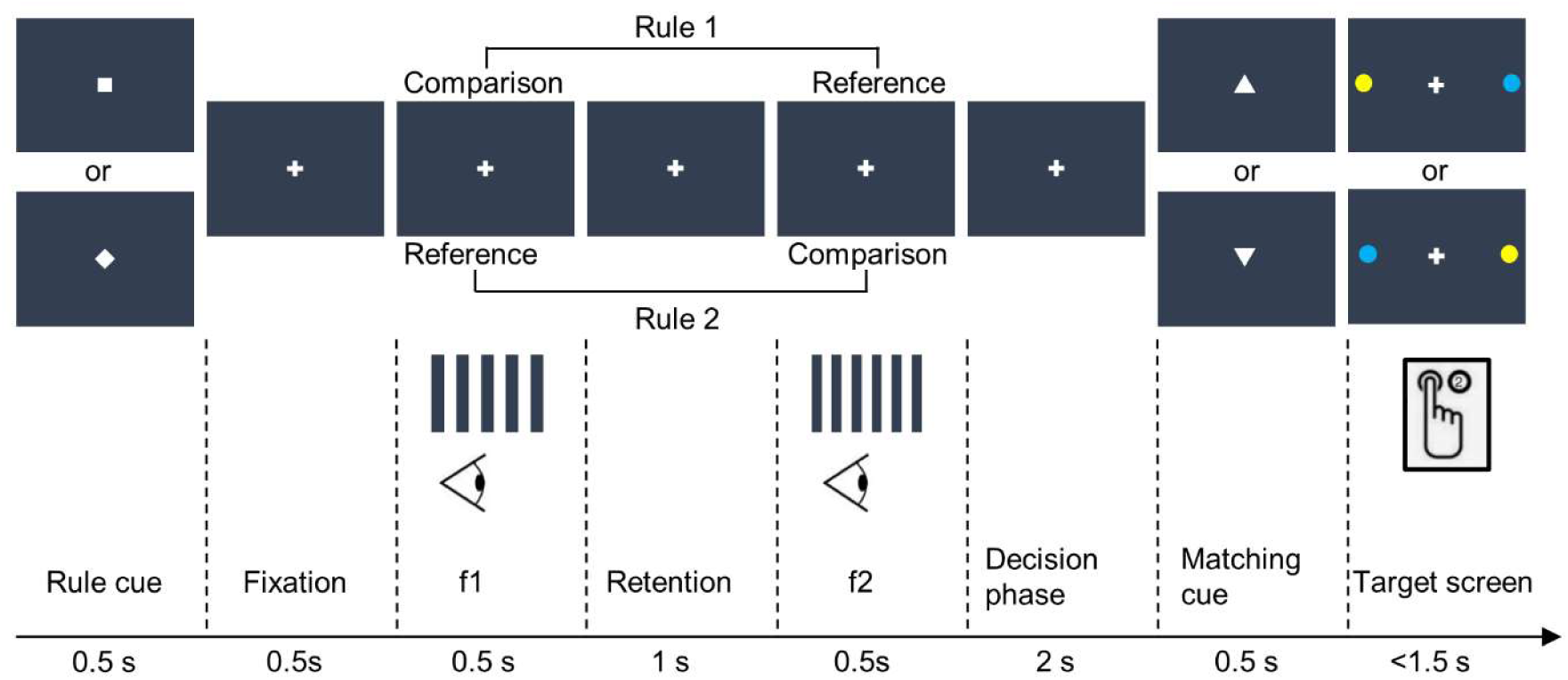
Experimental paradigm. An initial rule cue (square or diamond) indicated whether f1 would serve as the comparison and f2 as the reference stimulus (rule 1), or vice versa. Subsequently, participants had to decide whether the frequency of the comparison stimulus was higher or lower than the frequency of the reference stimulus. Following a short fixation period, two visual flicker stimuli with differing frequencies were sequentially presented. After the decision phase, one of two matching cues was presented. An upwards pointing triangle denoted that the frequency of the comparison stimulus was higher than the frequency of the reference stimulus while a downward pointing triangle represented a higher frequency of the comparison stimulus. Participants then had to decide whether their perception matches the matching cue and indicate their decision with a button press associated to one of two coloured disks on the following target screen.

Trials started with the presentation of one of two rule cues (i.e. Wu et al., 2019, 2021). Depending on its shape (square or diamond), the rule cue determined whether participants had to compare f1 against f2 or to compare f2 to f1 in half of the trials, respectively. The rule cue was introduced to decouple the decision (higher vs. lower) from potential effects of stimulus order (f1 > f2 vs f1 < f2). After a short fixation period, two consecutive visual flicker stimuli were presented on both sides of the participants periphery (5°, eccentricity) for 0.5 seconds, separated by a retention interval of one second. Following a two-second decision phase, a matching cue was presented, consisting of a triangle pointing either upwards or downwards. Participants had to compare their decision to the orientation of the matching cue (an upwards pointing triangle meaning “higher”, downwards meaning “lower”), to decide for “match” or “mismatch”. This procedure was introduced to render decisions independent from the motor response to avoid response-related confounds. The matching cue was independent from the true frequency difference and matches/mismatches were balanced within each run. Finally, participants reported the match or mismatch during the presentation of a target screen, displayed until the response but for a maximum of 1.5 seconds. The target screen comprised a central fixation cross and two coloured target disks (blue and yellow) in the periphery along the horizontal meridian (3°, eccentricity). The colour assignment for match or mismatch was balanced across participants. Depending on the location of the target, participants responded with a left or right button press, using their right-hand index or middle finger rendering the motor response independent of the perceptual decision. The target side was balanced within a run. Inter-trial intervals with a fixation period of varying durations (3,4,5 or 6 seconds) were administered between all trials.

During the fMRI session, the visual cues were projected onto a screen on the bore opening of the MR scanner. Participants viewed the visual displays through a mirror attached to the MR head coil from approximately 110 ± 2 cm. The cues were generated with MATLAB version 9.13 (The MathWorks, Inc, Natick, MA) using Psychtoolbox-3 (Brainard, 1997). Visual flicker stimuli were generated using a sine function with a fixed voltage amplitude of 10 V. In each trial, the sine waves were received and stored by a data acquisition card (NI-USB 6343; National Instruments Corporation, Austin, Texas, USA) and released upon a trigger signal to ensure precise timing. The visual flicker stimuli were generated with light emitting diodes (LEDs), transmitted through fibre-optic cables, and presented 10 cm to the left and right of a fixation cross on both sides of the screen. The LEDs illuminated above a threshold of 2.24 V such that the duty cycle of the flicker stimuli was approximately 43 %.

After training the task for 20-40 minutes, participants performed six experimental runs inside of the fMRI scanner on a separate day. A run lasted approximately 12.5 minutes and consisted of 64 trials with each of the eight frequency combinations being presented eight times per run. Each of the eight presentations contained a unique combination of rule cue, matching cue as well as target screen.

### Bayesian Model of Behavioural Data

To analyse the behavioural data and to estimate the individually perceived SFDs used for the fMRI analysis, we adopted a Bayesian model previously established for the vibrotactile DMTC task (Herding et al., 2016, 2019). Thereby, perceptual decisions (f1 > f2 or f1 < f2) are modelled by the integration of two subjective frequency distributions, which reflect noisy realizations of f1 and f2. To account for the Fechner’s law (Fechner, 1860) frequencies were log-transformed. In addition, the model accounts for the TOE (i.e. contraction bias), which is a commonly found bias in DMTC tasks (Ashourian & Loewenstein, 2011; Pennock et al., 2021; Saarela et al., 2023). The TOE refers to the phenomenon that the mnemonic representation of stimuli tends to regress towards the average of the stimulus set. The Bayesian inference model accounts for the TOE by combining the noisy distribution of the log-transformed f1 (likelihood distribution) with a distribution centred around the mean of a-priori observed log stimuli frequencies (prior distribution) to obtain a posterior distribution (f1’) which is shifted towards the mean of the overall stimulus set (Ashourian & Loewenstein, 2011; Boboeva et al., 2024). Thus, during the decision the subjective, mean-shifted posterior f1’ was compared to the noisy distribution of the log-transformed f2, leading to distinct performance patterns depending on stimulus order and f1 frequency (f2 is not retained and thus not combined with the prior distribution). A graphical illustration of the model is depicted in Fig. 2.

**Fig. 2.**
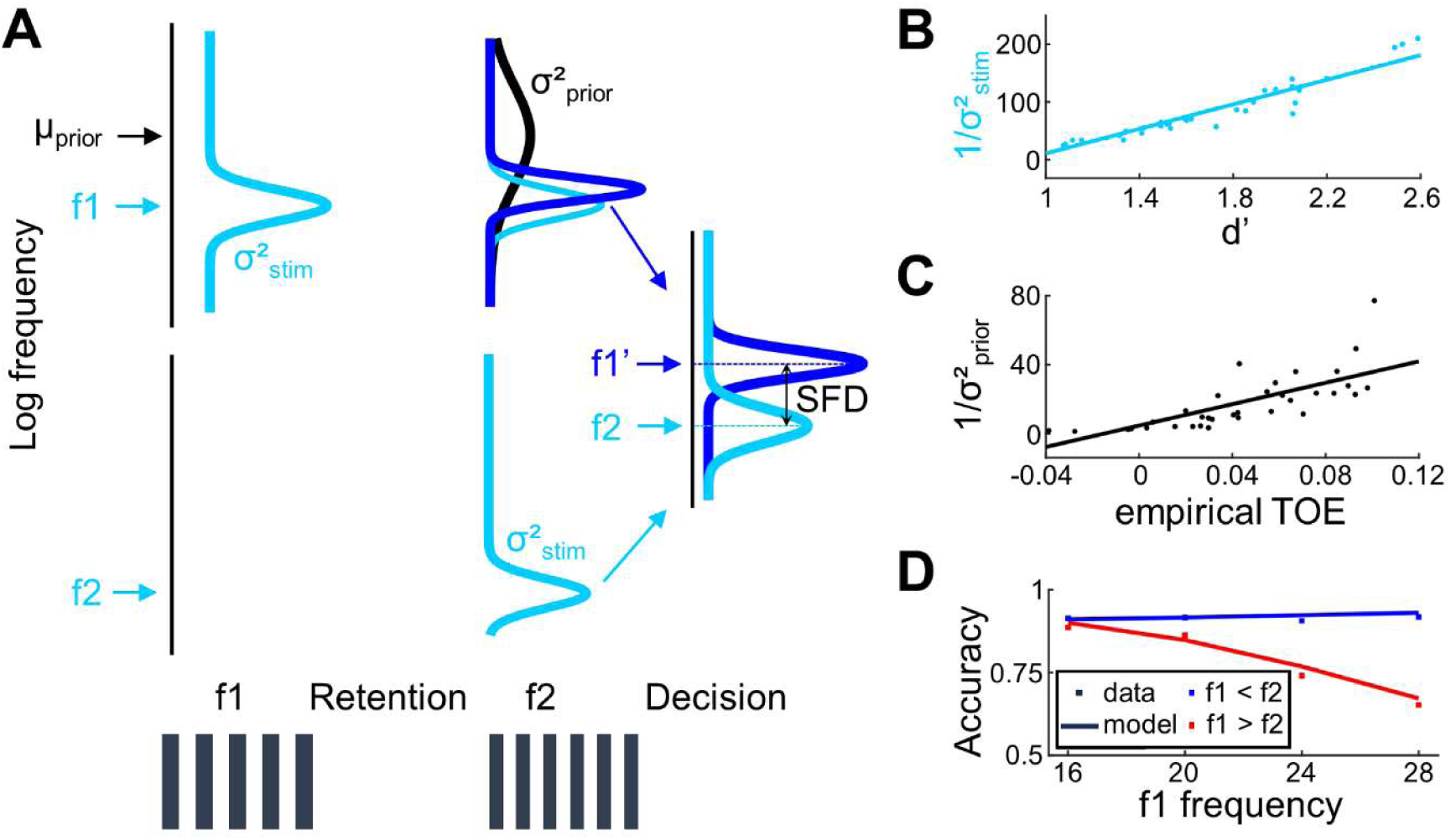
Graphical illustration of the Bayesian model. **A**. The three y-axes (black) depict perceived frequency values (log scale according to Fechner’s law). The top part of the graph illustrates how the representation of f1 develops as the task progresses. Upon presentation, f1 is represented as its likelihood function (cyan distribution). At the end of the retention period, f1 is combined with the prior distribution of the overall stimulus set (black distribution) to form f1’ (blue distribution; posterior distribution). In the decision, f1’ is compared to f2. The resulting SFD was used to model the decision strength within the fMRI decoding analysis. **B**. The scatter plot depicts the discriminability index (d’) plotted against the estimated stimulus precision (1/σ^2^_stim_). The discriminability index is calculated as d’ = z(hit rate) - z(false alarm rate). The hit rate equals to the proportion of f1 > f2 trials where participants correctly chose f1 > f2, and the false alarm rate is the proportion of f2 > f1 trials where participants erroneously chose f1 > f2. The strong positive correlation (ρ = 0.95, *p* < 0.001) reveals that the estimated precision serves as a good indicator of the participants accuracy. **C**. The scatter plot comparing empirical TOE vs. the prior precision also visualizes the strong positive correlation between the two measures (ρ = 0.754, *p* < 0.001), indicating that the prior precision is a good estimate for the TOE. **D**. Proportion of correct responses (squares) and model prediction of correct responses (lines) averaged across all participants. The model predictions closely resemble the observed data.

We included 3 free parameters in our model. (1) The variance of the likelihood distribution σ²_stim_ to model the individual’s perceptual precision (1/σ²_stim_) across all stimulus frequencies. A smaller σ²_stim_ indicated a less noisy perception of the stimulus frequencies. (2) The variance of the prior distribution σ²_prior_, which models the precision of the prior (1/σ²_prior_). A higher precision of the prior amplifies the influence of the stimulus mean and thus increases the TOE. (3) The response bias b indicates the participants tendency to prefer f1 > f2 over f1 < f2 or vice versa. These three model parameters were estimated for each participant individually by minimizing the variational free energy in a variational Bayes scheme, as implemented in the VBA toolbox (Daunizeau et al., 2014).

To quantify if the model with all three parameters (“full model”) provided the most accurate explanation for the behavioural data, we employed Bayesian model selection (BMS; Stephan et al., 2009) to compare it to a set of plausible alternative models. Our three comparison models included the “simple model” incorporating only σ²_stim_ as a free parameter, the “biased simple model” incorporating σ²_stim_ and b as free parameter, and the “unbiased TOE model” incorporating σ²_prior_ and σ²_stim_ as free parameters. In the applied BMS, models were regarded as random effects that could differ between subjects as implemented in the VBA toolbox (Daunizeau et al., 2014) and the variational free energy was used to select the best model (i.e. the model which occurs most frequently in the data set). By punishing for the inclusion of additional parameters, the variational free energy inherently accounts for model complexity and thereby avoids overfitting.

We used the individually estimated measures for the perceptual precision and the TOE to calculate SFDs of all 8 stimulus pairs by subtracting the posterior means of f1’ and f2. Since the means had to be subtracted in different orders, depending on the two varying comparison directions, a total of 16 SFD values (8 stimulus pairs * 2 comparison directions) were extracted for each participant. The SFDs were used to model the strength of the decision (i.e. the subjective decision values) in the subsequent fMRI decoding analysis.

### FMRI Data Acquisition and Preprocessing

Functional magnetic resonance imaging (fMRI) data was acquired on a 3 T Magnetom Prisma Fit Scanner (Siemens Healthcare GmbH, Erlangen, Germany) at the Center for Cognitive Neuroscience Berlin, using a 32-channel head coil. In each of the six experimental runs 378 functional, T2*-weighted volumes were acquired with a repetition time (TR) of 2000 ms, an echo time (TE) of 30 ms, an in-plane resolution of 64 x 64, a flip angle of 70° and a voxel size of 3 x 3 x 3 mm³. Furthermore, a T1-weighted image with 176 sagittal slices was acquired (TR = 1900 ms, TE = 2.52 ms, in-plane resolution: 256 x 256, voxel size: 1 x 1 x 1mm³).

### FMRI Analysis

Pre-processing and general linear model (GLM) analysis of the fMRI data was performed with SPM12 version v7388 (http://fil.ion.ucl.ac.uk/spm/). During the pre-processing, the functional images were slice-time corrected, realigned to the mean image and co-registered with the structural image.

To estimate voxel-wise decision-related activity patterns, we fitted a GLM with a 192 s high-pass filter to each participants functional data. Within each GLM, we estimated run-wise beta estimates during the decision phase for all voxels. The 16 regressors of interest included the 8 different stimulus pairs, each divided into the two comparison directions (comparing f1 against f2 vs comparing f2 against f1). The regressors were convolved with the hemodynamic response function at the onset of the decision phase. Further nuisance regressors included six movement parameters, the first five principal components to explain variance in white matter and cerebrospinal fluid signals respectively (Behzadi et al., 2007) and a run constant. Altogether 33 parameter estimates (16+6+5+5+1) were obtained for each run, resulting in a total of 198 regressors.

To identify brain regions with activation patterns being predictive of subjective decision values (i.e. SFDs), we applied an MVPA whole-brain searchlight approach for each participant. Voxel-wise activation patterns that exhibited a linear relationship with SFDs were obtained with a linear SVR using version 3.999F of The Decoding Toolbox (TDT; Hebart et al., 2015). A searchlight radius of 5 voxel was chosen as to implement a suitable payoff between sensitivity and specificity, where we confirmed the consistency of the main findings by additional control analyses with searchlight sizes of 4- and 6-voxel radius. Run-wise beta estimates of the GLM analysis were retrieved for all incorporated voxels of each searchlight location. The 16 SFD values from the Bayesian model were used as labels for an SVR. Then, a six-fold leave-one-run-out cross-validation procedure was applied, as implemented in TDT (Hebart et al., 2015). The resulting prediction accuracy maps comprised Fisher-z-transformed correlation coefficients at each searchlight location, depicting the correlation between true and predicted SFD labels. For the subsequent group-level analyses, the single-subject correlation maps were normalized to MNI space, resampled to a voxel size of 2 × 2 × 2 mm³ and spatially smoothed using a 3 mm full width at half maximum Gaussian filter. Correlation maps were then subjected to a one-sample t-test, to test for local brain activation patterns that showed significantly positive prediction accuracies of SFDs on the group level. All results are presented at p < 0.05 family-wise error corrected with a cluster extent threshold of k ≥ 50. To solidify the consistency of our findings, we additionally performed a simplified version of the decoding analysis by pooling the 16 SFDs into 4 pooled SFDs. The pooled SFDs were created by condensing the 4 lowest SFDs into one group, represented by their mean. This process was repeated for the next 4 higher SFDs and so on.

We conducted two additional decoding analysis to identify brain regions encoding information about the motor response and task rule. To this end, we implemented two separate GLMs with regressors modelling the motor response and task rule at the onset of the motor response or task rule, respectively. Similar to our main analysis, nuisance regressors were included to control for movement-related effects and variance in white matter and cerebrospinal fluid signals. Subsequently, two MVPA whole-brain searchlight approaches with a searchlight radius of 5 voxels were applied to the resulting beta images. A support vector machine (SVM) classifier using a six-fold leave-one-run-out cross-validation procedure was employed to find brain activity patterns with above-chance decoding accuracies for motor responses and task rules.

### Cross-Modal Analysis

One of our objectives was to investigate the neural underpinnings of subjective decision values across sensory modalities. Therefore, we performed the same SVR analysis, including the Bayesian modelling and SFDs in the decoding analysis, for the vibrotactile data obtained by Wu et al. (2021), who used the same study design and fMRI parameters. We kept all analysis parameters identical to our analysis. Applying the modelling of subjective decisions described above also to the data from Wu et al. (2021) allowed us to reintegrate the datasets of three participants, which were excluded in the original report due to poor behavioural performance. To reveal cross-modal decision-specific activation patterns, we entered the first level accuracy maps of both studies into flexible factorial second level model. Then, we computed a conjunction analysis to test the results against the global null hypothesis (Friston et al., 2005; Price & Friston, 1997).

## Results

### Behavioural Results

Participants performed the task with a mean accuracy of 84.9 % (SD: 7.2 %, range: 65.1-96.1 %) and an average response time of 519 ms (SD: 75 ms, range: 389-736 ms). Comparing behavioural performances to Wu et al. (2021), two-sample t-tests revealed no significant difference of mean accuracies (Δ_accuracies_ = −0.9 %, *p* = 0.625) while response times were slightly lower in our task (Δ_RTs_ = −46 ms, *p* = 0.045). Overall, the task difficulty was approximately equal between both studies. Comparing the effects of rule (compare f1 against f2 vs. f2 against f1), stimulus order (f1 > f2 vs. f1 < f2), and f1 frequency (16 Hz, 20 Hz, 24 Hz, and 28 Hz) as within-subject factors in a three-way repeated measure analysis of variance (ANOVA) revealed substantial performance differences across conditions (see Fig. 3). Similarly to Wu et al. (2021), there was no effect of task rule (F(1,35) = 0.585, *p* = 0.449) and a significant effect of stimulus order (F(1,35) = 68.588, *p* < 0.001). However, in contrast to Wu et al. (2021), this main effect was much stronger, and its direction was reversed, with a better performance in f1 < f2 trials (mean = 91.4 %) than in f1 > f2 trials (mean = 78.5 %). Similarly to Wu et al. (2021), the performance decreased with increasing f1 only in f1 > f2 trials while the performance in f1 < f2 trials remained the same throughout f1 frequencies, indicated by a significant main effect of f1 frequency (F(3,105) = 37.867, *p* < 0.001) as well as a significant interaction between stimulus order and f1 frequency (F(3,105) = 32.197, *p* < 0.001).

**Fig. 3.**
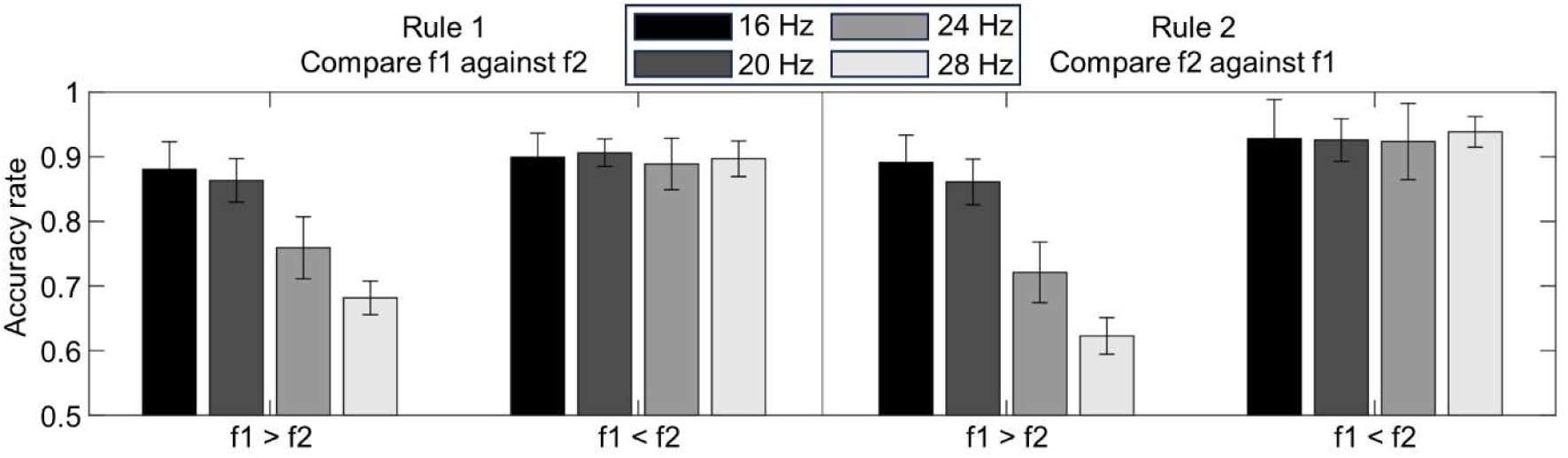
Behavioural results. The bar plots show the average performance across participants over all runs for different stimulus orders, rules and f1 frequencies. Error bars show the 95% confidence intervals (CIs).

We further tested whether potential biases in left and right motor responses within participants could have distorted the main decoding results. Note that the required motor responses were balanced through the task design. To that end, we computed Pearson chi-square tests comparing the distribution of the two conditions “higher” vs “lower” (indicating whether the true comparison frequency was higher or lower than the reference frequency) between left and right motor responses, for each participant. The results showed a significant difference for one participant (*p* = 0.045). In the remaining 35 participants, no significant difference was observed (all *p* > 0.1), suggesting that our main results were not influenced by imbalanced motor responses.

### Bayesian Model

The observed performance decrease for trials where f1 > f2 but not for trials where f1 < f2 (see ANOVA results and Fig. 3), suggests that the mere difference between f1 and f2 may reflect a combined influence of the TOE and the Fechner’s law. To account for these, we used the Bayesian modelling scheme to estimate subjective rather than objective frequency differences. The Bayesian model included the TOE as the σ²_prior_ (Mdn = 0.08), the precision as σ²_stim_ (Mdn = 0.015) as well as a general response bias *b* (Mdn = −0.056). The individually estimated parameters were closely related to empirical estimates. The estimated stimulus precision (1/σ²_stim_) was highly correlated with d’ values (ρ = 0.95, p < 0.001; Fig. 2B), while the empirical estimate of the TOE by Jamieson & Petrusic (1975) was sufficiently well described by prior precision (1/σ²_prior_; ρ = 0.754, *p* < 0.001; Fig. 2C). Furthermore, simulating choices based on the estimated model parameters predicted a performance pattern that closely resembles the observed data (see Fig. 2D). The individually estimated model parameters closely approximated performances also on the individual level.

To demonstrate the superiority of our proposed Bayesian model (“full model”) over a “null model”, which uses Fechner adjusted physical differences between f1 and f2 to quantify decisions, we calculated Bayes Factors comparing the free energies of the “full model” to the “null-model” on a single-subject level. This comparison revealed strong evidence in favour of the full model with Bayes Factors > 10 for 34 out of 36 participants (32 Bayesian *p*-values < 0.001, 2 Bayesian *p*-values < 0.05). Furthermore, with a random effects BMS framework, we compared the full model to three plausible comparison models with fewer parameters – the “simple model” (estimating only σ²_stim_ as a free parameter), the “biased simple model” (free parameters: σ²_stim_ and *b*) and the “unbiased TOE model” (free parameters: σ²_stim_ and σ²_prior_). The results of the BMS provided further evidence in support of the full model, showing an estimated frequency of 0.605 among the four compared models (exceedance probability = 0.976). Overall, the results indicate that a priori knowledge about the stimulus set (i.e. the TOE) substantially affects participants’ choices. Thus, participants most likely formed their decisions by comparing f2 to a version of f1 that regressed towards the mean of the stimulus set (i.e. f1’). Hence, we used the estimated SFDs as a proxy for subjective decision values in the fMRI analysis.

### FMRI Results

In the current study, our primary aim was to identify brain regions that convey information about subjective decision values, irrespective of stimulus order and motor responses. To accomplish this, we applied an MVPA, whole-brain searchlight approach during the 2-second decision phase. This approach allowed us to systematically test for brain regions that are predictive of the SFDs obtained from the Bayesian behavioural model. The results of the SVR, as depicted in Fig. 4, revealed local activation patterns with significant positive prediction accuracies (FWE corrected at *p* < 0.05 with a cluster extent threshold of k ≥ 50) in both hemispheres of the IPS as well as the PMd and in the left PMv. Hence, activation patterns in the PMd, the PMv and the PPC predicted subjective decision values irrespective of stimulus order and motor response. The cluster sizes of the identified brain regions are: 4132 voxels at the left IPS (peak voxel: [−28 −76 36]), 583 voxels at the right IPS (peak voxel: [34 −52 58]), 318 voxels at the left PMv (peak voxel: [−54 16 20]), 308 voxels at the right PMd (peak voxel: [30 8 42]), 128 voxels at the right IPL (peak voxel: [52, −26, 44]) and 52 voxels at the left PMd (peak voxel: [−20 6 54]). To further solidify the robustness of our findings, we repeated the SVR with different searchlight radii of 4 and 6 voxels, respectively. Both different searchlight radii reaffirmed the involvement of the same brain regions. Furthermore, we conducted a pooled decoding analysis with 4 instead of 16 distinct SFDs. The pooled analysis also identified similar predictive clusters than the main analysis. These included the left IPS (cluster size: 6885 voxels, peak voxel: [−28 −76 36]), the left PMv and PMd (cluster size: 1655 voxels, peak voxel: [−58 14 20]), the right PMd (cluster size: 685 voxels, peak voxel: [30 8 42]) and the left dorsolateral prefrontal cortex (cluster size: 80 voxels, peak voxel: [−36 38 2]).

**Fig. 4.**
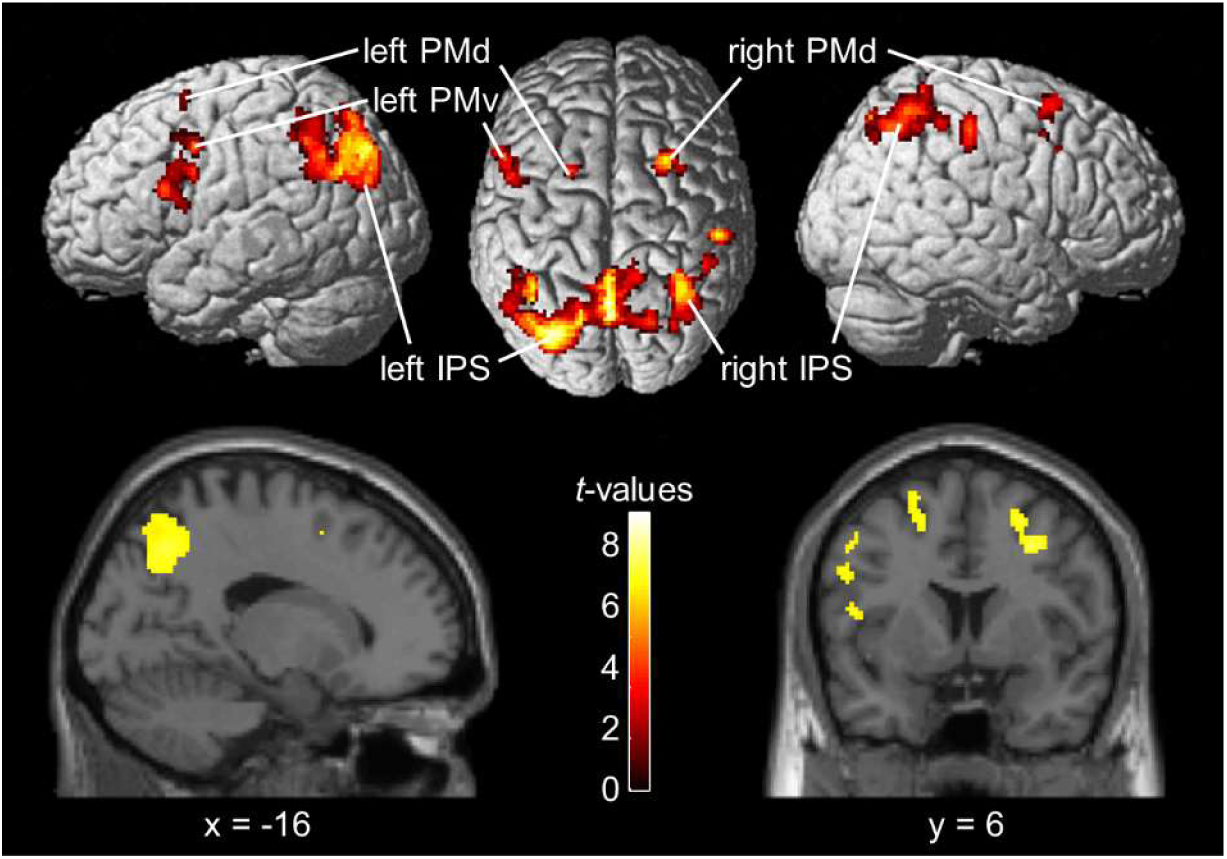
Results of the SVR approach, using SFDs as labels. The panel above depicts left, above and right view of the rendered standard brain with the projection of the brain regions predictive of subjective decision values, independent of stimulus order and motor response direction. The panel below depicts a medial and coronal view at the MNI coordinates x = −16 and y = 6, respectively. The SVR revealed activation pattern in the bilateral IPS and PMd as well as in the left PMv that were predictive of subjective decision values. Results are displayed FWE corrected at *p* < 0.05 with a cluster extent threshold of k ≥ 50.

To test for differences in prediction accuracy between hemispheres for both IPS and the PMd, single subject prediction accuracies were extracted from the peak voxel of each hemisphere and compared with two-sided paired t-tests. The t-test comparing left vs right IPS revealed a significantly higher decoding accuracy in the left IPS (t(35) = 2.728, *p* = 0.01). However, the t-test comparing left and right PMd did not indicate a significant difference between hemispheres (t(35) = −0.094, *p* = 0.926).

We additionally employed two SVM searchlight analyses to test for brain activation patterns with above-chance decoding accuracies of motor response and task rule during the decision period. Activation patterns selective for the motor response (left vs. right) were observed in the left motor cortex as well as in the bilateral occipital cortex. Activation patterns selective for the rule (compare f1 against f2 vs. f2 against f1) were observed in both hemispheres of the lateral occipital cortex, in the left IPS and in the left dorsolateral prefrontal cortex.

### Cross-Modal Analysis

The second aim of our study was to identify brain regions that carry information on subjective decision values across different sensory domains. To achieve this, we conducted a conjunction analysis against the global null hypothesis (Friston et al., 2005; Price & Friston, 1997) to compare our results to data of a vibrotactile study (Wu et al., 2021), which used an identical task design. Notably, Wu et al. (2021) employed a simpler decoding scheme, using a SVM to predict binary decisions. To ensure the comparability with our results, we re-analysed their dataset by applying a Bayesian model scheme as described above and the SVR. Subsequently, we tested the accuracy maps of both studies with a conjunction analysis to assess which brain regions could predict subjective decision values throughout both modalities. The conjunction analysis revealed positive prediction accuracies in several regions mostly surrounding the bilateral IPS and left PMd. The clusters included two separate clusters within the left IPS (cluster 1: peak voxel: [−14 −52 58], cluster size = 2568; cluster 2: peak voxel: [−28 −90 30], cluster size = 213), two separate clusters within the right IPS and IPL (cluster 1: peak voxel: [44 −36 52], cluster size = 1089; cluster 2: peak voxel: [30 8 42], cluster size = 329), a cluster in the left PMd and left PMv (peak voxel: [−26 8 52], cluster size = 1014), two clusters in the right PMv (cluster 1: peak voxel: [56 16 30], cluster size = 176; cluster 2: peak voxel: [44 36 −4], cluster size = 51) and a cluster in the right SPL (peak voxel: [14 −78 54], cluster size = 53). The results of the conjunction analysis are depicted in Fig. 5 (FWE corrected at *p* < 0.05 with a cluster extent threshold of k ≥ 50). Overall, the brain regions involved in encoding subjective decision values exhibited a great overlap across sensory domains, indicating that decisions are encoded within effector-specific brain regions, independently of the specific sensory modality.

**Fig. 5.**
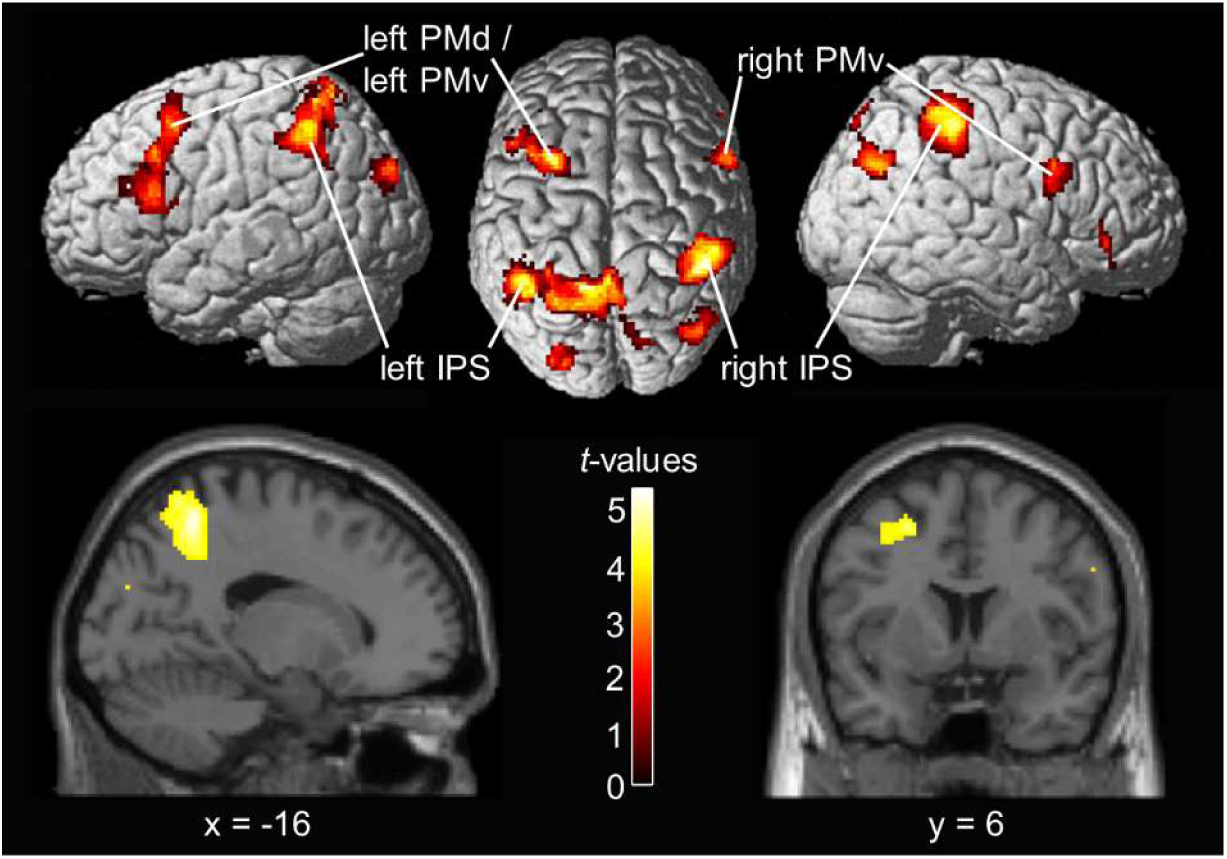
Results of the conjunction analysis against global null hypothesis. The panel above depicts the left, above and right view of the rendered standard brain with the projection of the brain regions predictive of subjective decision values across the visual and vibrotactile stimulus modality. The panel below depicts a medial and coronal view at the MNI coordinates x = −16 and y = 6, respectively. The conjunction confirmed positive prediction accuracies in the bilateral IPS, the left PMd and the left PMv. Furthermore, it revealed a positive prediction accuracy in the right PMv. Results are displayed FWE corrected at *p* < 0.05 with a cluster extent threshold of k ≥ 50.

## Discussion

In this fMRI study we aimed to test for effector-specific neural correlates of decisions in the visual domain that are independent from a specific motor response. Thus, we used a visual version of a DMTC task that rendered decisions independent from choice direction and response selection, with button-presses as a pre-specified effector. Furthermore, by employing Bayesian behavioural modelling alongside an SVR MVPA-searchlight approach, we were able to pinpoint brain regions predictive of subjective decision values. The SVR revealed positive prediction accuracies in the bilateral PMd, the left PMv and the bilateral IPS. These regions are highly similar to those observed in a similar vibrotactile DMTC study from Wu et al. (2021), suggesting a substantial cross-modal overlap between visual and vibrotactile sensory domain. To identify the cross-model neural correlates of subjective decision values, we applied the same modelling and decoding approach to their data and computed a conjunction analysis with the present data. The conjunction analysis revealed positive prediction accuracies in the left PMd, the bilateral PMv and the bilateral IPS. Hence, activation patterns in the PMd and the PPC consistently predicted subjective decision values not only regardless of stimulus order and motor response but also irrespective of stimulus modality. Thus, alongside previous findings by Wu et al. (2019, 2021), our results provide strong evidence that the effector-specific encoding of perceptual decisions is not only independent of task-related processes but that it also generalizes across sensory modalities.

### Modelling the Time-Order Effect Using Subjective Decision Values

Unlike previous DMTC studies, we focused on unravelling the neural correlates of subjective decision values rather than considering objective categorical decisions. Therefore, we employed a Bayesian inference model to individually compute SFDs for each stimulus pair, thereby avoiding assumptions of a constant perception of frequency differences across conditions. The SVR that we applied used the SFDs as labels that represented a measurement of subjective decision values. This approach allowed us to elaborate more fine-grained neural correlates of the decision. Prior EEG studies that employed the same Bayesian model approach suggested that EEG signals in both premotor (Herding et al., 2016) and posterior parietal regions (Herding et al., 2019) reflect subjective comparisons between two vibrotactile frequencies irrespective of choice accuracy. Our results corroborate these findings. Employing a Bayes optimal inference model to individually estimate subjective decision values enables a robust and realistic explanation of distinctive behavioural features in the given DMTC task. The applied model effectively captures the commonly observed TOE, while providing an explanation that preserves the optimality of participants’ behaviour (Ashourian & Loewenstein, 2011). Integrating such subjective biases could further refine our understanding of the TOE and consequently enhance our knowledge of the neural processes involved in perceptual decision-making.

### The Role of the Premotor Cortex in Perceptual Decision-Making

The premotor cortex has been found to play a crucial role in perceptual decisions by a series of primate vibrotactile studies which used arm movements or button presses as responses (reviewed in Romo & de Lafuente, 2013). The activity within premotor regions has been linked to different stages of the decision-making process, likely reflecting the integration of mnemonic processes and sensory inputs towards the formation of behavioural responses (Hernández et al., 2002; Wallis & Miller, 2003). Results from primate studies suggest that the PMd and PMv is involved in transforming sensory information into action through an evidence accumulation process (Cisek & Kalaska, 2005; Lemus et al., 2009; Romo et al., 2004; Wang et al., 2019). While the PMv has been associated to processes linked to action selection, the PMd has been more strongly associated with action preparation (Pardo-Vazquez et al., 2008; Pardo-Vázquez et al., 2011). However, recent primate research has supported the notion that the PMd is involved in more complex, associative processes beyond action preparation (Rossi-Pool et al., 2017). Similarly to Wu et al. (2021), our study decoupled the decision from task-related processes, notably motor responses and stimulus order, to identify the precise role of premotor cortices in perceptual decision-making. Both our study and the findings from Wu et al. (2021) indicate that the left PMd is involved in decision-making, irrespective of these task-related processes, supporting the notion that the PMd encodes categorical decisions. In addition to the original findings by Wu et al. (2021), we found positive prediction accuracies in the left PMv. Previous research has highlighted the important role of the PMv in perceptual decision-making (Pardo-Vázquez et al., 2011). Moreover, the PMv was predictive of subjective decisions in the conjunction analysis, suggesting a consistent contribution to the effector-specific representation that persists across sensory modalities. Furthermore, we found positive prediction accuracies in the right PMd in our main analysis and in the right PMv in the conjunction analysis. In contrast to the contralateral findings, these ipsilateral results are inconsistent between analyses and thus do not convincingly challenge the notion of a lateralized encoding of decision-related information in the premotor cortex. However, it remains possible that the more abstract effector-specific representations investigated in our study could be represented partially bilateral. Overall, our findings support the relevance of the premotor cortex in perceptual decision-making, even when participants have no knowledge about the upcoming motor response if the effector is pre-specified.

### The Role of the Posterior Parietal Cortex in Perceptual Decision-Making in Primates

A substantial body of primate studies investigating the neural correlates of perceptual decision-making has focused on the role of the PPC, with particular emphasis on LIP neurons (Gold & Shadlen, 2007). Most visual decision studies suggested that LIP neurons are predominantly involved in an effector-specific evidence accumulation process among competing saccade responses (Roitman & Shadlen, 2002; Shadlen & Newsome, 2001). However, almost all these studies used visual random dot motion tasks with saccades as response effectors, making it difficult to draw clear conclusions on the precise involvement of posterior parietal regions. Accordingly, recent primate studies have challenged this notion, showing that LIP neurons encoded perceptual decisions even in situations where the response was unpredictable during stimulus presentation (Shushruth et al., 2022) or disrupted (So & Shadlen, 2022). Freedman and colleagues conducted a series of studies employing a delayed match-to-category task with arbitrary categories of different random dot motion stimuli and manual arm movements as response modalities. Their results indicated that activity in LIP neurons reflect categorical decisions independent of specific effectors or stimulus modalities (Freedman & Assad, 2006; Swaminathan et al., 2013; Swaminathan & Freedman, 2012). Furthermore, by pharmacological inactivation, they demonstrated that LIP neurons have a causal role in evaluating task-relevant sensory stimuli that goes beyond the previously suggested primary function of merely representing motor responses (Zhou & Freedman, 2019). Interestingly, LIP inactivation impaired performance levels across different decision-making tasks regardless of the response modality used (Zhou et al., 2023). Overall, findings from primate studies suggest that neuronal activity in the PPC reflects decisions beyond a specific effector modality. Thus, despite its repeatedly demonstrated involvement in perceptual decision-making, the precise role of the PPC remains unclear.

### Sensorimotor Mapping in the Posterior Parietal Cortex

Similar to the abovementioned primate studies, human neuroimaging studies also challenge an exclusively effector-specific role of the PPC in perceptual decision-making. For instance, different regions of the PPC exhibited sustained activity throughout both arm reaching and saccadic responses, albeit with local variations in effector preference (Levy et al., 2007). Along these lines, a recent behavioural study on visual learning showed that training effects were only partially transferred between saccadic and reach responses. This partial learning transfer suggests that perceptual learning is neither entirely effector-specific nor completely effector-independent but rather entails a sensorimotor mapping from visual regions to effector-specific integrator regions. Given the PPC’s localization within the sensorimotor hierarchy as well as its overlapping encoding of multiple effectors, it emerges as a likely candidate region for a sensorimotor mapping across effectors (Ivanov et al., 2024). This notion was further supported by findings from the vibrotactile DMTC task by Wu et al. (2021), who showed that the IPS was predictive of binary decisions regardless of whether the effector was saccades or button-presses. Our results replicate these findings in a perceptual decision task in a different sensory modality, indicating that the IPS does not only represent decisions beyond the effector modality but also independent of the stimulus modality. Overall, this implies that the PPC conveys an abstract decision variable across multiple domains, with a varying degree of effector-specificity.

### Effector-Specific Representations Across Sensory Modalities

As described in the sections above, findings from primate studies overwhelmingly provide evidence for fully and partly effector-specific representations of visual decisions in premotor and posterior parietal regions. Conversely, when decoupling the decision from the motor response, human imaging studies suggest that decisions tend to be represented, either in a more abstract manner in posterior parietal and prefrontal brain regions (Filimon et al., 2013; Heekeren et al., 2008) or in brain areas associated with sensory specific stimulus features (Liu & Pleskac, 2011). In contrast, when decoupling decisions from the motor response in a vibrotactile DMTC task, Wu et al., (2019, 2021) demonstrated that activation patterns in effector-specific brain regions (in the FEF for saccadic responses and in the PMd for button press responses) were still predictive of categorical decisions. Comparing these findings to studies in the visual domain (e.g. Liu & Pleskac, 2011) indicates that there might be an interaction between response modality and stimulus modality. However, using a visual DMTC task with button presses as the effector modality, our results revealed activation patterns in highly similar effector-specific brain regions that also persisted in the cross-modal comparison between the visual and vibrotactile stimulus domains. Additional insight can be provided from the mouse model by Wu et al. (2020). They used optogenetics to inactivate premotor areas in an olfactory delayed match to sample task. Similar to our study, the mice knew the effector modality but were not able to plan a specific motor movement before the response. The disruption of neurons in the premotor cortex impaired task performance, suggesting that the relevance of premotor regions extend beyond the preparation of movements. Along with our results, it appears that the encoding of subjective decision values in effector-specific regions does not strictly constitute a mapping that is bound to a specific motor-movement. Rather, effector-specific brain regions appear to retain abstract information about stimulus identity and subsequently transforming them into motor responses.

### Flexible Representations of Perceptual Decision Processes

The findings from human studies seem to be somewhat incoherent, with some of them supporting an effector-specific representation of perceptual decisions while others suggest more abstract or stimulus-modality specific representations (Liu & Pleskac, 2011). However, upon further examination, this apparent discrepancy may be attributed to fundamental differences in analysis methods and task design. Firstly, our study employed a multivariate approach that aims towards uncovering nuanced local cortical activation patterns instead of focussing solely on average BOLD-signal. Consequently, distinct patterns that rely on both activation and deactivation may have been averaged out in analyses that use conventional fMRI analyses. Secondly, and perhaps more importantly, our study rendered decisions independent of a specific motor response while still preserving dependence on the effector itself. In contrast, prior studies that aimed to disentangle motor responses and decisions relied on abstract associations between decision and motor response, wherein participants were unaware of the effector until making their response. Overall, this implies that decisions are represented flexibly, depending on task demands. When a task does not pre-specify an effector, a decision between stimuli cannot be directly mapped to the respective effector-specific brain regions. Instead, information remains within brain areas associated with the sensory modality (Liu & Pleskac, 2011) or is transferred to a central, more abstract decision hub, e.g. in posterior parietal regions (O’Connell et al., 2012; Sandhaeger et al., 2023). On the other hand, if an effector is pre-specified, decisions are categorized as abstract motor intentions in effector-specific brain regions, facilitating the transformation into specific motor plans and actions upon presentation of the response mapping.

## Conclusion

In conclusion, our fMRI study suggests that if a response effector is pre-specified, local brain activation patterns encode subjective decision information in an effector-specific manner. Notably this encoding remains independent of stimulus order, motor response and even stimulus modality. It is conceivable that the PMd processes such effector-specific decisions signals, while the IPS adopts a more abstract representation of perceptual decisions which persists across different effectors. Moreover, our results indicate that activation patterns in these brain regions do not only represent categorical decision information but rather a more fine-grained representation of subjective decision values. Overall, our findings support the notion of a cross-modal, effector-specific representation of perceptual decisions if a response effector is pre-specified. Thereby, our study contributes to a deeper understanding of perceptual decision-making across a variety of sensory contexts.

## Data and code availability

The data that support the findings of this study are available on request to M.F.E. In accordance with EU’s General Data Protection Regulation and specifications in data protection section of the participants consent forms, we cannot share raw fMRI data.

## Author contributions

Marlon F. Esmeyer: Conceptualization, Data acquisition, Data curation, Formal analysis, Investigation, Methodology, Software, Validation, Visualization, Writing – original draft. Timo T. Schmidt: Conceptualization, Methodology, Software, Supervision, Validation, Visualization, Writing – review and editing. Felix Blankenburg: Conceptualization, Funding acquisition, Methodology, Project administration, Resources, Supervision, Validation, Writing – review and editing

